# Triad-LMF: A Hierarchical Low-Rank Multimodal Fusion Framework for Robust Cancer Subtype Classification Using Multi-Omics Data

**DOI:** 10.1101/2025.05.15.653805

**Authors:** Xingyue Tan, Xiran Chen, Renjie Tian, Qinyu Cai, Miaoyuan Jiang, Dongqiu Yang, Lei Zhang

**Affiliations:** School of Mathematics and Statistics, Chongqing Jiaotong University, Chongqing, 400070, China; Institute of Computer Vision and Traffic Image Understanding, School of Information Science and Engineering, Chongqing Jiaotong University, Chongqing, 400070, China; School of Life Sciences, Westlake University, Hangzhou,310030,China; Department of Bioinformatics, School of Basic Medical Sciences, Chongqing Medical University, Chongqing, 610101, 400016, China

**Keywords:** Cancer subtyping, Deep learning, supervised learning, low-rank multimodal fusion, hierarchy network

## Abstract

Cancer heterogeneity poses a major challenge for accurate molecular subtype classification. Conventional methods often fail to exploit complementary information across multiple omics modalities, leading to overfitting on high-dimensional data and limited representation of subtype heterogeneity. To address this, we propose Triad-LMF, a multiomics integration framework based on low-rank multimodal fusion to improve classification accuracy. Triad-LMF harmonizes heterogeneous omics data and integrates information through a two-stage hierarchical fusion strategy. Local Pairwise Fusion and Global Triadic Fusion are combined via the Two-Feature and Three-Way LMF modules, enabling a gradual transition from local modality interactions to global feature integration. Experimental results show that Triad-LMF consistently outperforms existing methods. UMAP visualization confirms enhanced subtype separability, and SHAP-based analysis highlights biologically meaningful features. Across independent datasets, Triad-LMF demonstrates strong generalization, offering a robust and interpretable framework for multiomics-driven cancer subtype classification.

## 1 Introduction

The high heterogeneity of cancer makes its molecular subtype classification a key cornerstone of precision medicine [1]. In recent years, the rapid development of multiomics data has provided unprecedented opportunities to uncover molecular mechanisms of cancer, identify subtype-specific biomarkers, and guide personalized therapy [2]. By integrating multimodal data such as gene expression, epigenetics, proteomics, and pathology images, researchers can more comprehensively depict the biological characteristics of cancer. However, the high dimensionality of multiomics data, the complex interactions between modalities,and data heterogeneity pose severe challenges to traditional computational methods. Therefore, developing novel integration frameworks that efficiently fuse these data is urgently needed to advance cancer subtype classification and precision medicine.

Supervised learning methods hold a significant position in cancer subtype classification models due to their ability to substantially improve classification Accuracy by leveraging labeled data. These methods mine subtype-specific patterns by integrating gene expression, mutation data, and other omics features, demonstrating excellent performance across cancer types such as breast cancer and colorectal cancer [3–5]. By optimizing model architectures and feature selection strategies, they enhance robustness against sparse or high-dimensional data, providing strong support for precise diagnosis and treatment. However, the limitations of existing methods cannot be overlooked: many adopt a one-time fusion strategy, making it challenging to capture dynamic interactions among modalities [5, 6]; handling high-dimensional data often leads to overfitting, especially pronounced in deep learning models [7, 8]. These shortcomings constrain the further application of multiomics data in precision medicine.

Based on the challenges above, we believe the fundamental issue of current models lies in the lack of hierarchical fusion mechanisms and explicit modeling of modality interactions, which limits the depth of biological pattern mining, while overfitting issues undermine model generalization. Therefore, this study proposes the Triad-LMF model, which introduces the following innovative designs to achieve breakthroughs in multiomics data integration and subtype classification:

- Triad-LMF first explicitly models modality-specific interactions through the Two-Feature LMF module, followed by high-level integration via the Three-way LMF module. This hierarchical design overcomes the limitations of traditional one-step fusion. Compared with existing methods, this strategy captures dynamic interactions between modalities through stepwise fusion, significantly improving classification performance.
- Low-rank decomposition techniques are introduced at all fusion stages. By enforcing low-rank constraints, the nonlinear modeling capability is enhanced while the number of parameters is significantly reduced, effectively alleviating overfitting issues caused by high-dimensional data and providing a feasible solution for large-scale multiomics data analysis.
- Triad-LMF prioritizes learning joint representations of specific modality pairs through the Two-Feature LMF module, generating interpretable interaction features that improve model robustness. It also offers biological insights into the contribution of modality pairs to subtype classification, deepening the exploration compared to existing methods and providing new perspectives for identifying subtype-specific biomarkers.

## 2 Related Work

### 2.1 Unsupervised Learning

Unsupervised learning methods mine potential subtypes through clustering or dimensionality reduction. For example, [9]proposed a kernel fusion approach that integrates gene expression, miRNA expression, and isoform data to construct similarity kernel matrices, thereby efficiently clustering breast cancer and lung cancer subtypes. This method uses kernel functions to capture nonlinear relationships between modalities and demonstrates high subtype discrimination capability on the TCGA dataset. However, its simple fusion strategy makes it difficult to model hierarchical interactions between modalities, limiting the analysis of complex biological patterns. [10] developed a multiview spectral clustering method (MRF-MSC) that fuses multiomics data through multi-smooth representations and combines spectral clustering to uncover subtype structures, showing excellent performance across multiple cancer types in the TCGA dataset.[11] proposed SMMSN (Self-supervised Multi-fusion Strategy Network), which integrates multiomics data through graph convolutional networks and stacked autoencoders using dual self-supervision for label-free cancer subtype discovery, demonstrating significant clinical relevance in survival analysis and case studies across eight cancer datasets.

### 2.2 Supervised Learning

To achieve higher classification Accuracy, supervised learning methods leverage labeled data to improve cancer subtype classification performance significantly. BCDForest by [3] optimizes subtype classification of small-scale gene expression data through multi-granularity scanning and enhances model robustness to sparse data via random forests and multi-scale feature selection strategies; however, its performance is limited when generalized to large-scale multiomics datasets. DeepCSD by [4] employs a minimalist feedforward neural network combined with gene expression and mutation data to identify colorectal cancer subtypes efficiently. It benefits from low computational complexity but has limited capacity to capture complex biological features.

Li et al. [5, 12, 13] utilize graph convolutional networks (GCNs) to integrate multiomics data by constructing intermodality relation graphs, demonstrating excellent performance in breast cancer and pancancer classification. These works exploit graph neural networks to mine topological relationships between modalities but suffer from high model complexity, easily leading to overfitting. [6, 14, 15] optimize classification and biomarker identification through GCNs and graph attention networks (GATs), combining attention mechanisms to prioritize key features, further enhancing cancer subtype classification Accuracy and interpretability. [16] perform multi-class classification using co-expression modules, mining gene co-expression patterns to strengthen the model’s biological relevance, but remains insufficient in modality fusion.[8] propose a robust GCN framework that stabilizes high-dimensional data through regularization optimization, achieving excellent performance in pan-cancer subtype classification. [17] model molecular mechanisms via gene interaction networks, significantly improving subtype classification Accuracy. Although these supervised methods excel in Accuracy, their one-time fusion strategies struggle to model dynamic interactions between modalities, and handling high-dimensional data often leads to overfitting issues.

### 2.3 Semi-Supervised Learning

In scenarios with limited annotated data, semi-supervised learning offers a compromise solution. DeepType by [1] integrates supervised classification and unsupervised clustering, effectively leveraging the limited labeled information in the TCGA dataset through joint optimization of labeled and unlabeled data, demonstrating excellent breast cancer subtype classification performance. [2] mined nonlinear features using a semi-supervised model that combines autoencoders and regularization techniques, enhancing the model’s generalization ability on small sample datasets. [18] utilized multiomics autoencoders to predict breast cancer subtypes, uncovering latent features through nonlinear dimensionality reduction and improving model robustness.[19] optimized classification performance by incorporating consistency regularization, enhancing classification Accuracy by constraining model consistency on unlabeled data. [20] modeled gene mutation data via mutation networks combined with graph neural networks to optimize classification, showing potential in pancancer subtype classification. However, these fusion methods still exhibit insufficient modeling of modality interactions, making it difficult to capture the complex relationships in multiomics data comprehensively.

## 3 Methods

This paper proposes a deep learning framework called Triad-LMF based on Low-rank Multimodal Fusion (LMF). Inspired by [21], this method efficiently fuses heterogeneous omics data through modality-specific low-rank factors, overcoming the limitations of traditional tensor fusion methods and generating a unified low-dimensional representation for downstream cancer subtype identification. Figure 1 shows the framework of Triad-LMF. We describe each block in more detail in the following.

**Figure. 1.**
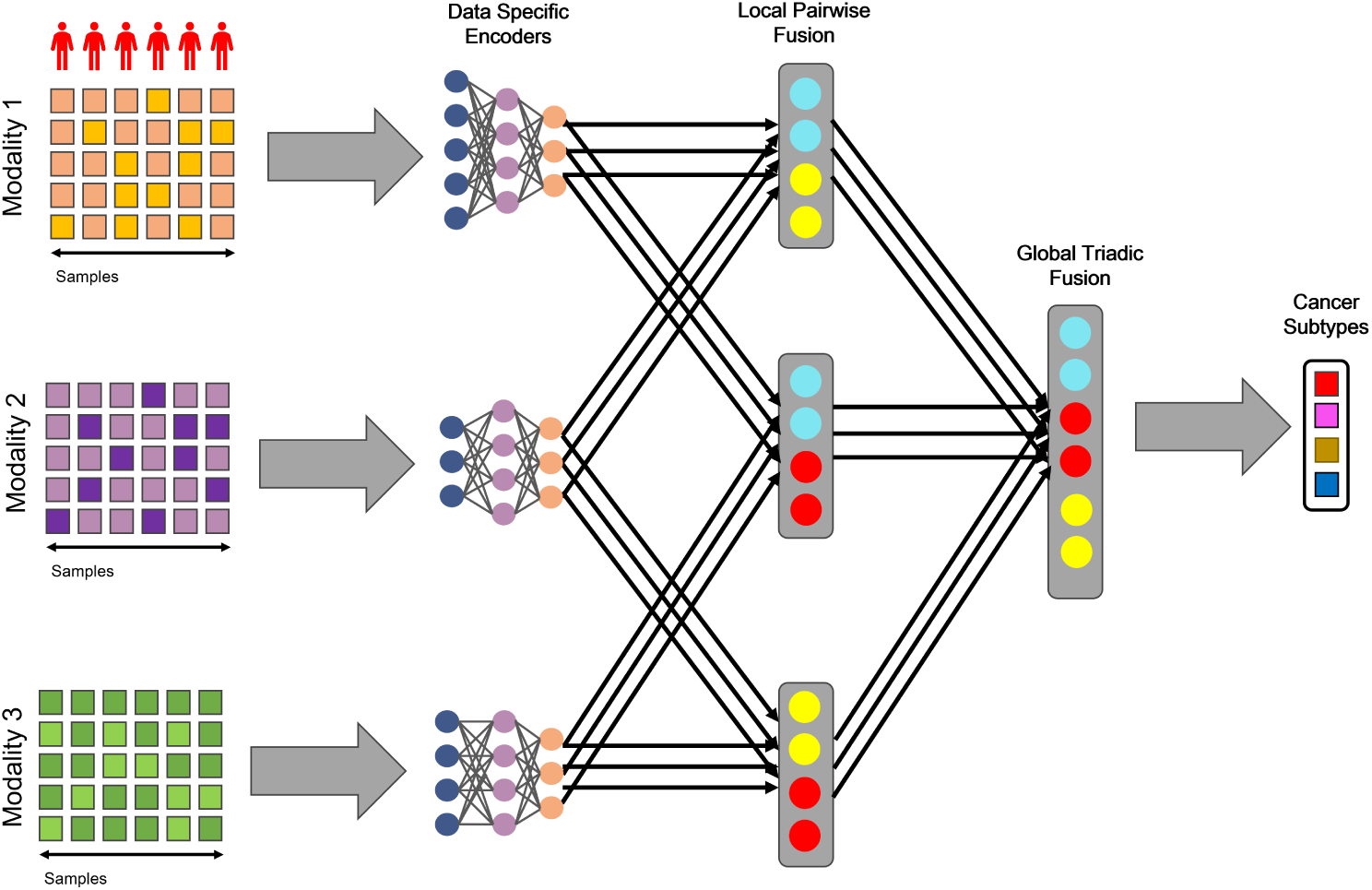
The framework of Triad-LMF

### 3.1 Modility-Specific Feature Extractor Encoder

Multiomics data typically exhibit high dimensionality, heterogeneity, and noise interference, which pose significant challenges for feature extraction. For example, the RNA-seq data in the BRCA dataset contains 16,383 features, where RPPA includes only 223 features. This discrepancy in dimensionality and data noise may obscure underlying biological patterns. To address this, we designed a Modality-Specific Embedding Subnetwork.

Given a set of cancer multiomics data *X* = {*X*_1_*, X*_2_*, . . . , X_N_* },*X_i_* ∈ *R^B×di^ , i* = 1, 2*, . . . , N* , where *N* is the number of datasets, *X_i_* is the input data of modality *i*, with *d_i_* as the input dimension of modality. To extract the hidden representations *X*^^^*_i_* ∈ *R^B×d^*^^^*^i^* of nonlinear features for each modality. Each omics is assigned with Modality-specific Encoder *X*^^^*_i_* = *f_i_*(*X_i_*).

For each omics modality, we employ an encoder network composed of three fully connected layers to extract high-level latent representations. The input features are first normalized using Batch Normalization to stabilize training and mitigate internal covariate shift. A dropout layer is then applied to reduce the risk of overfitting by randomly deactivating neurons during training.

Each encoder layer performs a linear transformation followed by a ReLU activation to introduce non-linearity and enhance feature expressiveness. Specifically, given an input feature vector, the encoding process can be expressed as:

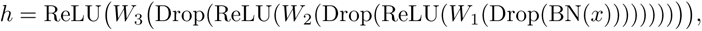

where *W_i_* denotes the weight matrix of the *i*-th linear layer, BN(·) represents Batch Normalization, and Drop(·) denotes the Dropout operation.

### 3.2 Low-rank Multimodal Fusion

Integrating representations from multiple modalities to capture cross-modal interaction information in multiomics data analysis is crucial for achieving precise cancer subtype classification. However, traditional multimodal fusion methods often result in exponential growth in computational complexity when dealing with high-dimensional data, which is impractical for real-world applications. We designed Low-rank Multimodal Fusion modules that significantly reduce computational complexity by introducing modality-specific low-rank factors to address this issue. We propose two variants for this purpose:

- Local Pairwise Fusion for pairwise modality fusion
- Global Triadic Fusion is used to integrate all modalities.

#### 3.2.1 Local Pairwise Fusion

Given the representation of two modalities *X*_1_ ∈ *R^B×h^*^1^ and *X*_2_ ∈ *R^B×h^*^2^ , respectively, the fused output representation generated by the two feature LMF is *Z* ∈ *R^B×r^*, where *r* is the rank parameter. To introduce a bias term and enhance the flexibility of the representation, we first extend the hidden representations, as defined in Eqs. (1) and (2):

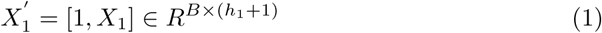

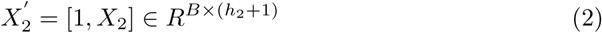

where 1 is an all-one vector.

The fusion process uses modality specific low-rank factors *F*_1_ ∈ *R^r×^*^(^*^h^*^1 +1)^*^×dy^* and *F*_2_ ∈ *R^r×^*^(^*^h^*^2 +1)^*^×dy^* , where *d_y_* is the output dimension. The intermediate fusion tensor is calculated as following Eqs. (3) and (4):

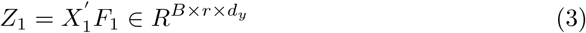

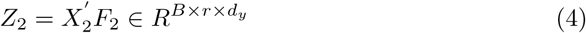

where *Z*_1_ and *Z*_2_ ∈ *R^B×r×dy^* are modality-specific intermediate representations. Then, cross-modal interaction characteristics are captured through element-wise multiplication, as formulated in Eqs.(5):

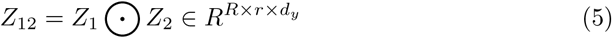

where *Z*_12_ is the joint tensor. The final output *Z* ∈ *R^B×r^* is obtained by extracting the first dimension and transposing, as formulated in Eqs. (6):

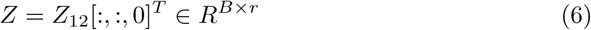

Therefore, the output of Local Pairwise Fusion generates a compact feature vector of dimension r for each sample, where *r* is the low-rank parameter reflecting the degree of compression of the fused representation. This representation not only efficiently preserves cross-modal correlations but also provides high-quality intermediate features for subsequent Global Triadic Fusion multimodal integration.

#### 3.2.2 Global Triadic Fusion

Given three representations *H*_1_,*H*_2_,*H*_3_ produced by the Modility-Specific Feature Extractor Encoder, The final output representation of the Global Triadic Fusion is *O* ∈ *R^B×dy^* , with *d_y_* as the output dimension, which we set to the number of specific cancer subtype categories.

It is worth noting that the Global Triadic Fusion does not directly fuse the original hidden representations produced by the Modality-Specific Feature Extractor Encoder. Instead, it is built upon the results of three bimodal low-rank fusions described in Section.(3.2.1). This hierarchical design is first used to extract local interactions between modalities through pairwise fusion, and then applied to integrate global features via trimodal fusion. This enhances the model’s robustness and expressive power in scenarios with modality imbalance.

The implementation of Global Triadic Fusion relies on a preceding pairwise fusion step. The specific process is as follows: Before Global Triadic Fusion, three Local Pairwise Fusions respectively fuse modality pairs to generate intermediate representations as shown in Eqs. (7)–(9):

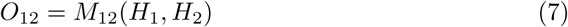

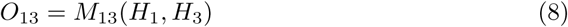

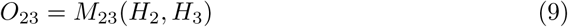

Where *M_ij_* denotes the Local Pairwise Fusion, which outputs *O*_12_*, O*_13_*, O*_23_ ∈ *B^B×r^*, *r* is the rank parameter, and each output is a compact feature vector of dimension *r* corresponding to each sample.

Take the three pairs of fused intermediate representations *O*_12_*, O*_13_*, O*_23_ as the input of the Global Triadic Fusion. Similarly, extend its hidden representation as shown in Eqs. (10)–(12):

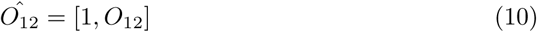

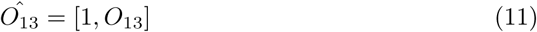

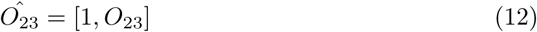

where 1 ∈ *R^B×^*^1^ is an all-one vector, and the extended representations are denoted Define low-rank factors for each intermediate representation as modality-specific factors *F*_12_ ∈ *R^r^ ^×^*^(^*^h^*^1 +1)^*^×dy^* , *F*_13_ ∈ *R^r^ ^×^*^(^*^h^*^2 +1)^*^×dy^* , *F*_23_ ∈ *R^r ×^*^(^*^h^*^3 +1)^*^×dy^* , where *r* is the rank parameter of the three-modality fusion, and *d_y_* is the final output dimension. Then, we use low-rank factors to transform the expanded intermediate representation, as shown in Eqs. (13)–(15):

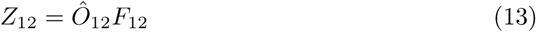

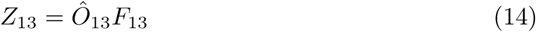

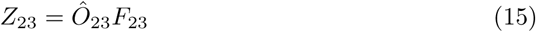

where *Z*_12_*, Z*_13_*, Z*_23_ ∈ *R^B×r^ ^×dy^* represents pairs of intermediate tensors. Next, integrate the three intermediate tensors through element-wise multiplication, as shown in Eqs. (16) :

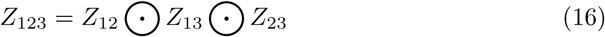

Where *Z*_123_ ∈ *R^B×r^ ^×dy^* is the joint tensor that captures the global interaction information among all modalities. The joint tensor generates the final output through a weighted sum and bias calculation, as shown in Eqs. (17):

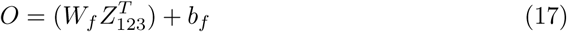

Where *W_f_* ∈ *R*^1*×*^*^r^* is the fusion weight matrix, *b_f_* ∈ *R*^1*×*^*^dy^* is the bias vector, and the reshaped output *O* ∈ *R^B×dy^* is the feature vector corresponding to each sample with dimension *d_y_*.

Compared to Local Pairwise Fusion, Global Triadic Fusion performs hierarchical fusion, which not only preserves local interaction characteristics but also achieves efficient integration of global information.

### 3.3 Hierarchical Fusion Integrator

In the Triad-LMF framework, the fusion of multiomics data is realized through a hierarchical structure to efficiently integrate heterogeneous information and generate a unified representation suitable for cancer subtype classification. The Hierarchical Fusion Integrator, a core component of this layered design, integrates Local Pairwise Fusion and Global Triadic Fusion step-by-step via the hierarchy, transitioning from local modality interactions to global information integration.

Given the imput multimodal omics data *X*_1_*, X*_2_ and *X*_3_, its corresponding hidden representations are denoted as *H*_1_ ∈ *R^B×h^*^1^ , *H*_2_ ∈ *R^B×h^*^2^ and *H*_3_ ∈ *R^B×h^*^3^ , which are generated by the Modality-Specific Embedding Subnetwork. Here, *B* represents the batch size, and *h_m_*denotes the hidden dimension of each modality. The Hierarchical Fusion Integrator employs a two-stage strategy:

- Two-Feature LMF module: Three Local Pairwise Fusion modules are applied to modality pairs, producing intermediate representations that capture local inter-modal interactions.
- Three-way LMF module: A Global Triadic Fusion module integrates the above intermediate representations to generate the final output, which can directly support cancer subtype classification tasks.

The Hierarchical Fusion Integrator optimizes the contributions of sparse modalities and balances the representation weights across modalities. Its performance will be validated in subsequent experiments.

## 4 Experiments and Results

### 4.1 Dataset

We applied the proposed model to classification tasks on three different datasets. The first dataset is breast invasive carcinoma (BRCA), which includes CNV, mRNA, RPPA, and PAM50 subtype labels (including Her2, LumA, and LumB subtypes), sourced from the literature [5]. The second dataset is used for molecular subtype classification of STAD (including CIN, MSI, GS, and EBV subtypes), provided by the literature [17]. Both datasets contain a sufficient number of samples and a reasonable number of subtypes. Additionally, for BRCA cancer subtype classification, we introduced a third dataset: the PAM50 subtype classification of breast invasive carcinoma (BRCA 2) from TCGA. Compared to the BRCA dataset, which covers broader molecular levels, BRCA 2 focuses more on RNA regulatory mechanisms. All datasets are summarized in Table 1.

**Table 1.**
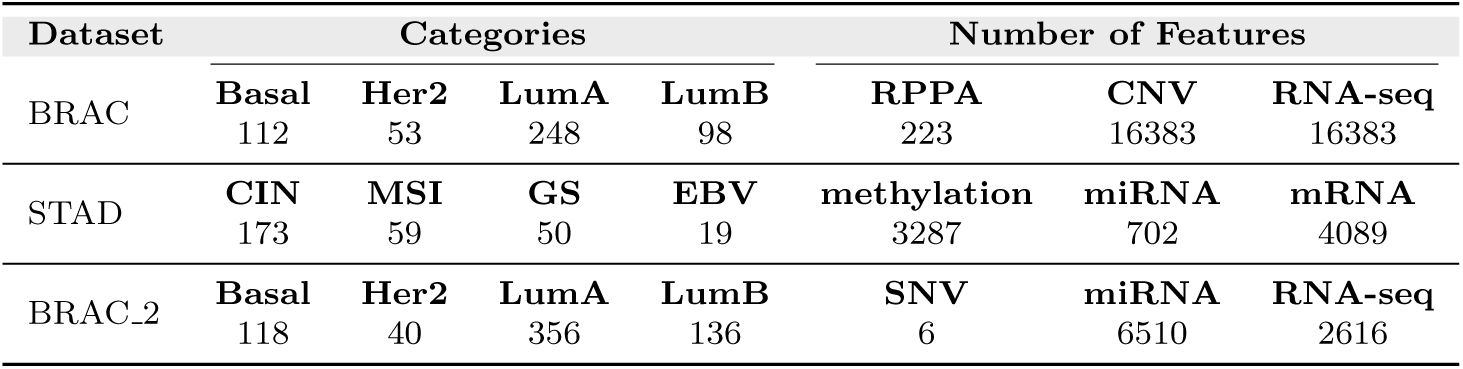
Description of BRAC, STAD, BRAC 2 dataset

| Dataset | Categories |  |  |  | Number of Features |  |  |
| --- | --- | --- | --- | --- | --- | --- | --- |
| BRAC | **Basal**<br>112 | **Her2**<br>53 | **LumA**<br>248 | **LumB**<br>98 | **RPPA**<br>223 | **CNV**<br>16383 | **RNA-seq**<br>16383 |
| STAD | **CIN**<br>173 | **MSI**<br>59 | **GS**<br>50 | **EBV**<br>19 | **methylation**<br>3287 | **miRNA**<br>702 | **mRNA**<br>4089 |
| BRAC_2 | **Basal**<br>118 | **Her2**<br>40 | **LumA**<br>356 | **LumB**<br>136 | **SNV**<br>6 | **miRNA**<br>6510 | **RNA-seq**<br>2616 |

It is worth noting that in the BRCA 2 dataset, the number of features in the SNV modality is only 6, showing a significant imbalance compared to miRNA (6510 features) and RNA-seq (2616 features). Such extreme differences in feature dimensions across modalities are common in multiomics data. This significant imbalance among modalities may weaken the model’s ability to capture sparse modalities when using traditional multimodal fusion methods (such as feature concatenation or tensor outer product), as high-dimensional ones can overshadow the contributions of low-dimensional modalities. The Triad-LMF model effectively addresses this challenge through its unique design, and its efficiency and robustness in imbalanced modality scenarios will be discussed in detail in Section 3. We preprocessed the data before applying the Triad-LMF model to ensure quality and consistency. First, patient samples with more than 10% missing information in any modality were excluded from the dataset. This threshold was chosen based on standard practices reported in the literature [8]while retaining sufficient samples to support statistical analysis. Second, features with more than 10% missing values across all patient samples were removed to avoid impairing the model’s feature extraction capabilities due to incomplete information. These preprocessing steps transformed the multimodal data into a high-quality set, laying a solid foundation for modality-specific embedding and multimodal fusion.

### 4.2 Evaluation Metrics

To comprehensively quantify Triad-LMF’s performance in cancer subtype multiclass classification tasks, we selected the following standard evaluation metrics.

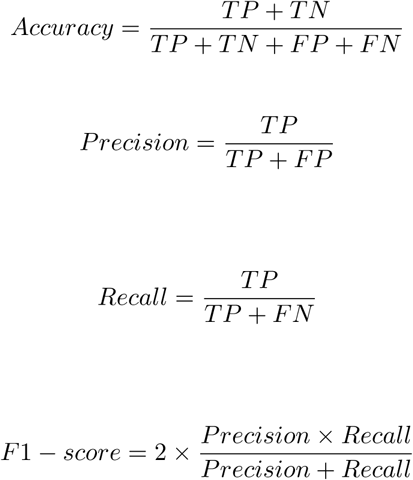

Where *TP* refers to samples that actually belong to a certain class and are correctly predicted as that class, while *TN* refers to samples that the model correctly identifies as not belonging to the target class. For multiclass tasks, any class other than the positive class is defined as negative. *FP* refers to samples the model incorrectly predicts as belonging to the target class, and *FN* refers to samples that the model incorrectly predicts as not belonging to the target class. *Accuracy*, *Precision*, *Recall*, and *F* 1−*score* are the most commonly used classification performance metrics based on the aforementioned *TP* , *TN* , *FP* , and *FN* . Considering the potential class imbalance in the dataset, we primarily report *F* 1 − *score* and *Accuracy* to evaluate model performance comprehensively. To further ensure the reliability of the results, all performance comparisons are validated for significance using the Wilcoxon rank-sum test.

### 4.3 Experimental Setup

The implementation of Triad-LMF is based on the PyTorch framework, and training and testing were conducted on an NVIDIA GeForce RTX 3060 GPU. To ensure the model’s stability and generalization capability, we performed a systematic hyperparameter optimization on the validation set through grid search, focusing on key parameters such as the learning rate, local fusion rank, global fusion rank, and dropout rate. Table 2 lists the candidate value ranges for the hyperparameters.

**Table 2.** Triad-LMF Hyperparameter Optimization Range

| Hyperparameter | Candidate value range |
| --- | --- |
| Learning Rate | $1e-3, 5e-4, 1e-4, 5e-5$ |
| Batch Size | 32, 64, 128 |
| Hidden Layer | 64, 128, 256 |
| $r$ | 2, 4, 8 |
| $r'$ | 4, 8, 16 |
| Weight Decay | $1e-4, 1e-5, 1e-6$ |
| Dropout | 0.1, 0.3, 0.5 |

Triad-LMF uses the cross-entropy loss function and minimizes the loss through the Adam optimizer. To prevent overfitting, we introduce an early stopping mechanism that terminates training when the validation loss does not decrease for 10 consecutive epochs. Additionally, it should be noted that Dropout is not mentioned in the table; we adjust the corresponding Dropout rates according to the characteristics of each modality to balance regularization effects. All experiments are repeated five times, reporting the metrics’ mean and standard deviation to ensure the results’ robustness and reproducibility. As shown in Table 3, the final selected hyperparameter combination achieves excellent classification performance on the validation set.

**Table 3.** Key Experimental Setup Parameters

| Hyperparameter | Value | Hyperparameter | Value |
| --- | --- | --- | --- |
| Framework | PyTorch | Weight Decay | $1e-5$ |
| Optimizer | Adam | $H_m$ | 128 |
| Learning Rate | $1e-4$ | $r$ | 4 |
| Batch Size | 64 | $r'$ | 8 |
| Training Epochs | 100 | Early Stopping $\delta$ | 10 |
| Weight Decay | $1e-5$ | Early Stopping Patience | 10 |
| Hardware Platform | NVIDIA GeForce RTX 3060 |  |  |

During training, we apply batch normalization to the preprocessed features *X_m_* to reduce input feature covariate shift. Moreover, we need to avoid overfitting in high-dimensional feature space since our datasets have a relatively small sample size. We applied Dropout regularization during training

### 4.4 Triad-LMF Outperforms Existing Methods for Comparative Cancer Subtype Diagnosis

To rigorously evaluate the performance of Triad-LMF, we compared it against a series of baseline methods and related advanced multiomics fusion approaches on the BRCA and STAD datasets. The baseline methods were constructed as follows:

- Naive multimodal fusion: directly concatenating feature vectors from all modalities, then inputting the concatenated high-dimensional features into traditional machine learning classifiers for training.
- Leading multiomics fusion methods: Triad-LMF was further compared with MOGONET [6], DeepMo[22], and MoGCN[5].

The same training-testing data splits were used across all comparative methods to ensure fairness in all experiments. Performance evaluation uniformly used the metrics defined in Section 3.1. The experimental results are shown in Table 4 and Figure 2. On both the BRCA and STAD datasets, Triad-LMF consistently outperformed all baseline and comparative methods on key evaluation metrics. On the BRCA dataset, Triad-LMF achieved an F1 score of 0.8613, while the best-performing baseline method, RF, scored 0.8435. Similarly, as shown in Table 5 for the STAD dataset, Triad-LMF reached an Accuracy of 0.8761, surpassing MOGONET’s 0.8197 and DeepMo’s 0.8462. These results demonstrate that our proposed hierarchical fusion strategy can effectively integrate information from different omics data types, thereby significantly improving the Accuracy of cancer subtype classification.

**Table 4.** Comparison of Triad-LMF Diagnostic Methods for Cancer Subtypes on BRCA Dataset

| Method | Accuracy | Precision | Recall | F1 |
| --- | --- | --- | --- | --- |
| DT | $0.8350 \pm 0.021$ | $0.7500 \pm 0.030$ | $0.7730 \pm 0.028$ | $0.7614 \pm 0.027$ |
| KNN | $0.6117 \pm 0.035$ | $0.3960 \pm 0.045$ | $0.3382 \pm 0.040$ | $0.3608 \pm 0.042$ |
| GNB | $0.5728 \pm 0.038$ | $0.3233 \pm 0.048$ | $0.3441 \pm 0.045$ | $0.3336 \pm 0.046$ |
| RF | $0.8830 \pm 0.015$ | $0.8710 \pm 0.018$ | $0.8174 \pm 0.020$ | $0.8435 \pm 0.019$ |
| SVM | $0.8447 \pm 0.018$ | $0.8379 \pm 0.020$ | $0.7773 \pm 0.022$ | $0.8068 \pm 0.021$ |
| MoGCN | $0.8002 \pm 0.016$ | $0.7919 \pm 0.034$ | $0.8015 \pm 0.010$ | $0.7967 \pm 0.018$ |
| DeepMo | $0.8626 \pm 0.025$ | $0.8079 \pm 0.017$ | $0.7871 \pm 0.010$ | $0.7871 \pm 0.010$ |
| MOGONET | $0.8738 \pm 0.016$ | $0.8837 \pm 0.017$ | $0.7644 \pm 0.019$ | $0.8196 \pm 0.018$ |
| **Triad-LMF** | **$0.8947 \pm 0.012$** | **$0.8950 \pm 0.013$** | **$0.8299 \pm 0.014$** | **$0.8613 \pm 0.013$** |

**Figure. 2.**
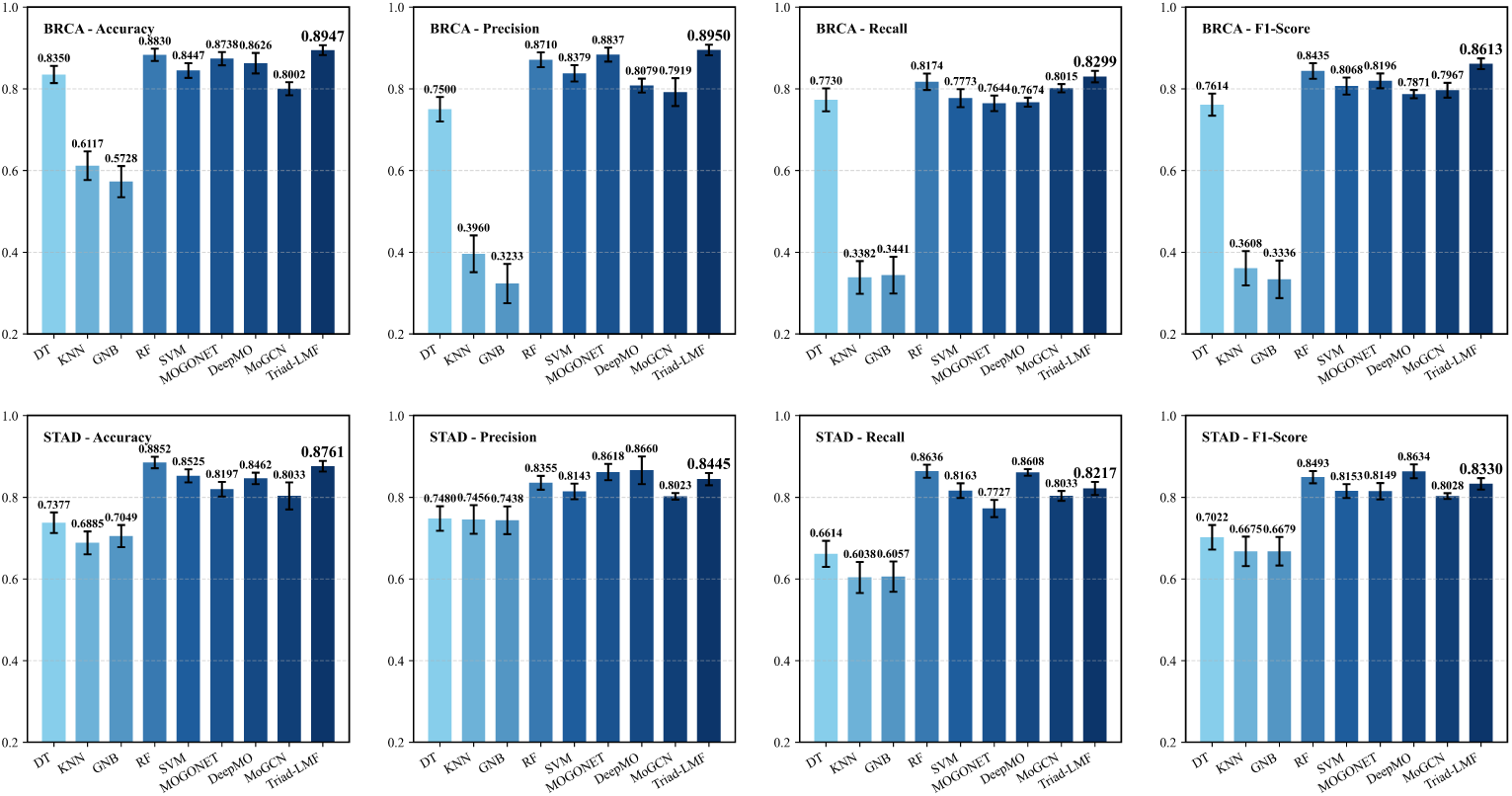
Triad-LMF outperforms existing cancer subtype diagnosis methods in standard supervised learning comparisons

**Table 5.** Comparison of Triad-LMF Diagnostic Methods for Cancer Subtypes on STAD Dataset

| Method | Accuracy | Precision | Recall | F1 |
| --- | --- | --- | --- | --- |
| DT | $0.7377 \pm 0.025$ | $0.7480 \pm 0.030$ | $0.6614 \pm 0.032$ | $0.7022 \pm 0.030$ |
| KNN | $0.6885 \pm 0.028$ | $0.7456 \pm 0.035$ | $0.6038 \pm 0.038$ | $0.6675 \pm 0.036$ |
| GNB | $0.7049 \pm 0.027$ | $0.7438 \pm 0.034$ | $0.6057 \pm 0.037$ | $0.6679 \pm 0.035$ |
| RF | $0.8852 \pm 0.014$ | $0.8355 \pm 0.017$ | $0.8636 \pm 0.016$ | $0.8493 \pm 0.015$ |
| SVM | $0.8525 \pm 0.016$ | $0.8143 \pm 0.019$ | $0.8163 \pm 0.018$ | $0.8153 \pm 0.017$ |
| MoGCN | $0.8033 \pm 0.033$ | $0.8023 \pm 0.008$ | $0.8033 \pm 0.012$ | $0.8028 \pm 0.007$ |
| DeepMo | $0.8462 \pm 0.014$ | $0.8660 \pm 0.034$ | $0.8608 \pm 0.008$ | $0.8634 \pm 0.017$ |
| MOGONET | $0.8197 \pm 0.018$ | $0.8618 \pm 0.020$ | $0.7727 \pm 0.021$ | $0.8149 \pm 0.020$ |
| **Triad-LMF** | **$0.8761 \pm 0.013$** | **$0.8445 \pm 0.015$** | **$0.8217 \pm 0.016$** | **$0.8330 \pm 0.014$** |

### 4.5 Revealing the Latent Space Structure of Triad-LMF through UMAP

To qualitatively analyze the Triad-LMF model’s ability to learn discriminative data representations during the hierarchical fusion process, we employed UMAP technology to reveal the model’s progressive refinement of cancer subtype discriminatory information through two-dimensional visualization. UMAP is a nonlinear dimensionality reduction method that effectively preserves both local and global topological structures of high-dimensional data, making it suitable for evaluating the discriminability of latent space representations[23]. Specifically, we extracted embedding vectors from four key stages of Triad-LMF’s hierarchical fusion architecture: three Local Pairwise Fusion stages and one Global Triadic Fusion stage. Based on test samples from the BRCA and STAD datasets, these vectors reflect the model’s representational capability for cancer subtype classification at different fusion stages.

We applied UMAP to project the high-dimensional embedding vectors into a two-dimensional space to assess samples’ clustering characteristics and intercluster separability in the latent space. As shown in Figure 3, during the Local Pairwise Fusion stages, the model already demonstrated preliminary subtype differentiation ability, with samples of the same subtype forming loose clusters and cluster boundaries exhibiting some level of separability. However, the final latent space representation after Global Triadic Fusion significantly outperformed the earlier stages, evidenced by tighter subtype clusters and more precise boundaries between clusters. These visualizations reveal the mechanism by which Triad-LMF progressively enhances representational discriminability through modal synergy, validating the effectiveness of its hierarchical fusion architecture in integrating multi-modal information.

**Figure. 3.**
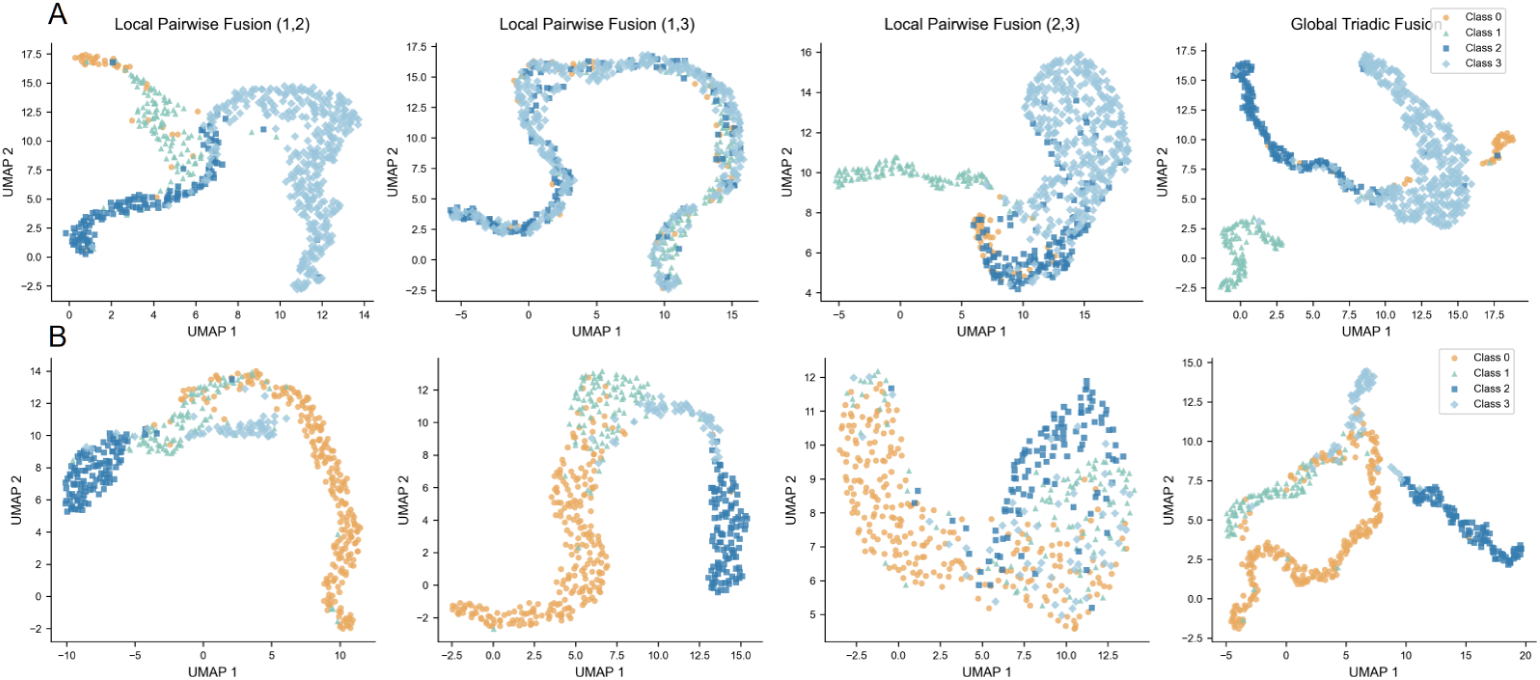
UMAP Visualization of STAD (A) and BRCA (B)

### 4.6 Ablation study

To quantitatively evaluate the contributions of each key component in the Triad-LMF framework—notably the Local Pairwise Fusion and Global Triadic Fusion—to the overall model performance, we designed and conducted a series of ablation experiments using the BRCA dataset. These experiments investigated the impact on performance metrics by sequentially removing certain components of Triad-LMF or replacing specific parts of the framework. The training loss dynamics are shown in Figure 4(A), and the final classification performance comparison is presented in Figure 4(B). The proposed Triad-LMF framework achieves the best performance in terms of the ACC metric, confirming its overall effectiveness. In contrast, removing any Local Pairwise Fusion module from Triad-LMF causes a significant drop in Accuracy, indicating that pairwise cross-modal interactions are crucial for enhancing classification performance and that the model effectively extracts valuable discriminative features from the fusion of modal pairs. Notably, when the Global Triadic Fusion is removed and replaced with simple concatenation, the model’s performance falls short of the full Triad-LMF and underperforms compared to the baseline model. This result highlights that simply obtaining pairwise fused features is insufficient; the subsequently designed low-rank tri-modal fusion mechanism is essential for effectively integrating information from all three pairwise interactions and capturing higher-order complex relationships rather than merely summing the information. Through systematic ablation experiments, we confirm that each core component of the Triad-LMF model contributes positively to the final performance. Removing any single component leads to a performance decline. This fully validates the rationality and effectiveness of our proposed hierarchical, two-stage fusion architecture, demonstrating that Triad-LMF can fully leverage both pairwise cross-modal interactions and higher-order tri-modal integrated information to achieve superior cancer subtype classification performance.

**Figure. 4.**
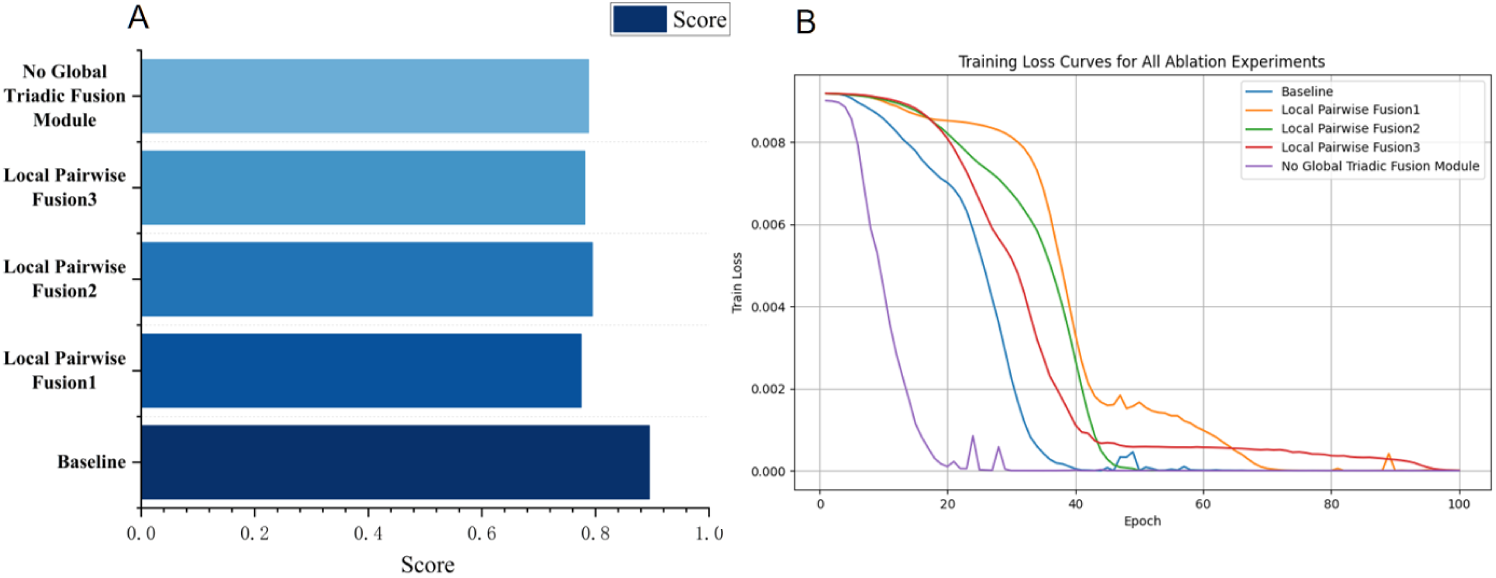
(A) BRCA ablation experiment Accuracy bar chart (B) BRCA ablation experiment loss function line chart (Local Pairwise Fusion1: removed the fusion of modalities 1 and 2 from the baseline. Local Pairwise Fusion2: removed the fusion of modalities 2 and 3 from the baseline. Local Pairwise Fusion3: removed the fusion of modalities 2 and 3 from the baseline. No Global Triadic Fusion: removed Global Triadic Fusion from the baseline, replaced with simple concatenation.)

### 4.7 Feature Importance Analysis and Biological Interpretation based on SHAP

To explain the cancer subtype classification mechanism of Triad-LMF and explore its biological significance, we employed the SHapley Additive exPlanations (SHAP) method[24]for a systematic analysis of feature importance in Triad-LMF. SHAP is a model interpretability tool based on game theory that quantifies the marginal contribution of each input feature to the model output, providing both global and local assessments of feature importance. We calculated SHAP values for all features in the test set of the BRCA dataset, with a focus on the features with the highest overall contributions and those that have a significant impact on the classification of specific cancer subtypes. Using SHAP’s TreeExplainer, we analyzed the prediction results of the Triad-LMF model, computed SHAP values for each feature, and ranked the features based on their mean absolute SHAP values. Table 6 lists the top 60 ranked features, encompassing key proteins, phosphorylation sites, and other biomarkers. These features play important roles in the molecular mechanisms of breast cancer, reflecting the model’s sensitivity to biologically relevant signals. Additionally, as shown in Figure 5(C), we present the sensitivity of the four BRCA subtype categories to the significant features.

**Table 6.** Top 60 Features Identified by Triad-LMF on the BRCA Dataset Based on SHAP

| Rank | Feature | Rank | Feature | Rank | Feature |
| --- | --- | --- | --- | --- | --- |
| 1 | S6 | 21 | EGFR_pY1173 | 41 | PDK1 |
| 2 | ERALPHA | 22 | S6_pS235S236 | 42 | PTEN |
| 3 | PR | 23 | SRC_pY527 | 43 | CYCLINE1 |
| 4 | P27_pT198 | 24 | G6PD7 | 44 | MYH11 |
| 5 | BCL2 | 25 | CYCLINB1 | 45 | GATA3 |
| 6 | P90RSK_pT359S363 | 26 | DVL3 | 46 | TFRC |
| 7 | CAVEOLIN1 | 27 | TUBERIN_pT1462 | 47 | PI3KP110ALPHA |
| 8 | EEF2 | 28 | ERALPHA_pS118 | 48 | ANNEXIN1 |
| 9 | EGFR | 29 | JAK2 | 49 | MAPK_pT202Y204 |
| 10 | IRS1 | 30 | LCK | 50 | BIM |
| 11 | SYK | 31 | AMPKALPHA_pT172 | 51 | CASPASE3 |
| 12 | CASPASE7CLEAVEDD198 | 32 | ANNEXINVII | 52 | SCD1 |
| 13 | PAXILLIN | 33 | EGFR_pY1068 | 53 | AMPKALPHA |
| 14 | HER3 | 34 | XRCC1 | 54 | RICTOR |
| 15 | ASNS | 35 | RBM15 | 55 | MYOSINIIA_pS1943 |
| 16 | RICTOR_pT1135 | 36 | IGFBP2 | 56 | GATA6 |
| 17 | MYOSINIIA | 37 | AR | 57 | HER2 |
| 18 | ACETYLATUBULINLYS40 | 38 | PARPCLEAVED | 58 | RAB11 |
| 19 | HER2_pY1248 | 39 | PDL1 | 59 | PCADHERIN |
| 20 | GSK3ALPHABETA | 40 | STAT3_pY705 | 60 | CHK2 |

**Figure. 5.**
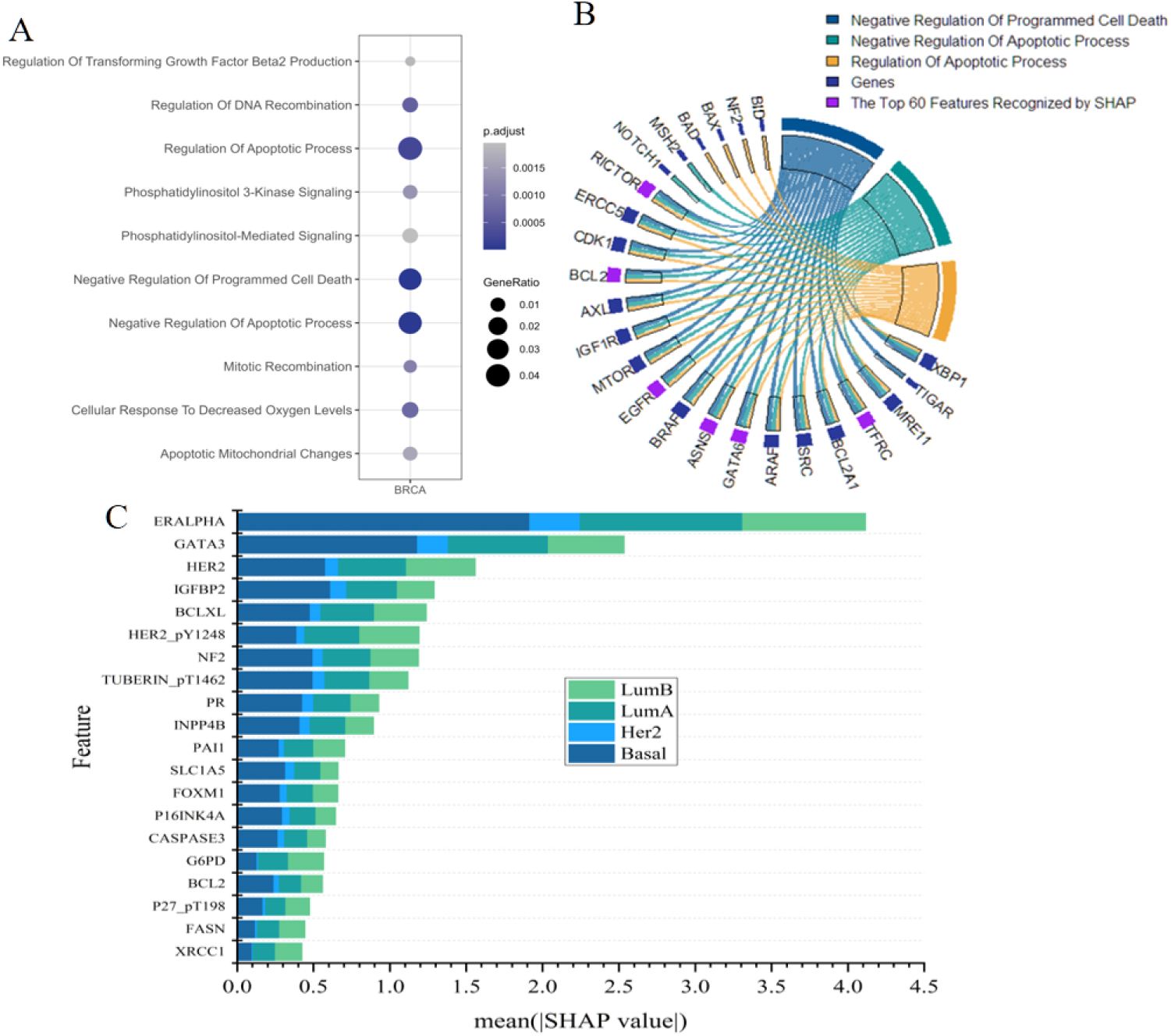
(A) GO functional enrichment analysis results of the top 60 features. (B) Average absolute SHAP values of the top 60 features. (C) Sensitivity of four BRCA dataset categories to significant features.

As shown in Table 6, the top-ranked features show the precision of the Triad-LMF model in capturing the key nodes of breast cancer biology. Ribosomal protein S6 is a core downstream effector of the mTOR signaling pathway, regulating protein synthesis and cell proliferation. Its significant ranking suggests that Triad-LMF is highly sensitive to mTOR-driven tumor proliferation. Aberrant activation of the mTOR signaling pathway is widespread in breast cancer and is closely associated with tumor progression and treatment resistance[25]. The primacy of S6 in the SHAP analysis is consistent with the clinical efficacy of mTOR inhibitors (e.g., everolimus), which studies have shown to significantly improve the prognosis of patients with hormone receptor-positive breast cancer [26, 27].ERALPHA and PR are signature molecules of the LumA and LumB subtypes that regulate hormone-dependent gene expression and tumor cell proliferation. The high ranking reflects the model’s prioritized focus on hormonal signaling pathways, which is highly consistent with breast cancer molecular typing studies [28].ERALPHA expression level is a key predictor of the efficacy of endocrine therapies (e.g., tamoxifen), and PR expression further refines prognostic assessment of the Luminal subtype[29, 30]. The ability of Triad-LMF to recognize both underscores its Precision in the classification of hormone receptor-positive breast cancer.P27 pT198 and BCL2 are involved in cell cycle regulation and anti-apoptotic mechanisms, respectively.P27, through the phosphorylation site T198 regulates nucleoplasmic localization and cell cycle progression, and its high ranking suggests that the model focuses on cell cycle dysregulation in breast cancer progression[31]. It has been shown that P27 phosphorylation correlates with poor prognosis, reflecting the active state of tumor proliferation [32].BCL2, as an anti-apoptotic protein, inhibits programmed cell death and promotes tumor survival. It is often highly expressed, especially in Luminal subtypes. The prominent ranking of BCL2 in the SHAP analysis suggests that the model effectively captures the tumor survival mechanisms, consistent with its role in breast cancer prognosis and treatment response [33]. eGFR pY1068, HER2 pY1248, and MAPK pT202Y204 mark the activation status of EGFR, HER2, and MAPK/ERK signaling pathways, respectively. These features are particularly prominent in HER2+ subtypes, driving cell proliferation and survival.HER2 pY1248 is a key target for targeted therapies (e.g., trastuzumab), whereas MAPK pT202Y204 is associated with tumor invasion and metastasis [34]. The high ranking of EGFR pY1068 is further suggestive of the model’s identification of the EGFR signaling pathway in breast cancer heterogeneity [35]. The outstanding contribution of these features suggests that the model Triad-LMF accurately captures the dynamics of the signaling network, consistent with the complexity of breast cancer molecular mechanisms [36]. In addition, PI3KP110ALPHA, PTEN, GATA3, and PDL1 regulate the PI3K/AKT pathway, differentiation characteristics of Luminal subtypes, and immune escape mechanisms in TNBC, respectively.PI3KP110ALPHA and PTEN co-regulate the PI3K signaling pathway, which is closely associated with tumor proliferation and treatment resistance [37].GATA3, as a transcription factor, is associated with the hyper differentiation characteristics of the Lum subtype, and its ranking reflects the model’s focus on subtype-specific features [38].PDL1, on the other hand, is a key molecule for immune escape in TNBC, which is correlated with the efficacy of immune checkpoint inhibitors, suggesting that the model integrates the tumor microcircuits with the tumor micro-circuits—correlates, suggesting that the model integrates tumor microenvironmental features [39]. Identifying these features further validates the model’s ability to integrate signal transduction, molecular differentiation, and immune regulation.

The top 60 features identified by SHAP are highly consistent with the known molecular mechanisms of breast cancer, encompassing mTOR, hormone receptors, PI3K/AKT, EGFR/HER2, and immune-related pathways. This validates the Triad-LMF model’s biological interpretability and highlights its potential application value in Precision medicine, providing an important basis for biomarker validation and targeted therapy research. Next, we conducted a Gene Ontology (GO) on the top 60 high-ranking features, focusing on the Biological Process category. The results showed significant enrichment (*p <* 0.05) in key pathways, including (1) negative regulation of programmed cell death involving BCL2, CASPASE7CLEAVEDD198, CASPASE3, and BIM, underscoring tumor anti-apoptotic mechanisms closely related to breast cancer cell survival and therapy resistance [33, 40]; (2) regulation of DNA recombination, including XRCC1 and CHK2, reflecting the role of DNA repair and genome stability in BRCA1/2 mutant subtypes [41, 42]; (3) phosphatidylinositol-mediated signaling, covering IRS1, PDK1, and PI3KP110ALPHA, pointing to the PI3K/AKT/mTOR pathway, which is associated with tumor proliferation and resistance to targeted therapies [26, 37]. These pathways are highly consistent with the molecular mechanisms of breast cancer, for example, the PI3K/AKT pathway driving cell survival [43] and DNA repair influencing genome instability subtypes [36]. Figure 5(B) visually illustrates the associations between these pathways and genes, while Figure 5(C) further quantifies pathway significance and gene counts. We confirmed that the Triad-LMF model can achieve interpretable subtype classification by capturing key biological processes.

### 4.8 Validating Effectiveness using other BRCA datasets

To evaluate the robustness and generalization ability of the Triad-LMF framework, we applied it to the BRCA 2 dataset. This dataset features the same cancer types and subtypes as BRCA but utilizes a different combination of omics modalities. We trained and evaluated the Triad-LMF model on the BRCA 2 dataset using the same hyperparameter settings applied to the original BRCA dataset.

Triad-LMF also demonstrated strong performance on the BRCA 2 dataset, achieving an accuracy of 0.9221, an F1 score of 0.9150, a precision of 0.9082, and a recall of 0.9219. As shown in Figure 6, both the training and testing accuracy and loss curves remain closely aligned throughout the learning process, indicating stable convergence and minimal overfitting. These results are comparable to those obtained on the BRCA dataset, confirming consistent performance across both datasets. This indicates that the proposed Triad-LMF framework is not limited to specific combinations of omics data types but effectively adapts to different modalities while maintaining robust classification performance for the same biological subtypes. Overall, these findings further support the universality and robustness of the proposed hierarchical fusion strategy for multiomics cancer data integration.

**Figure. 6.**
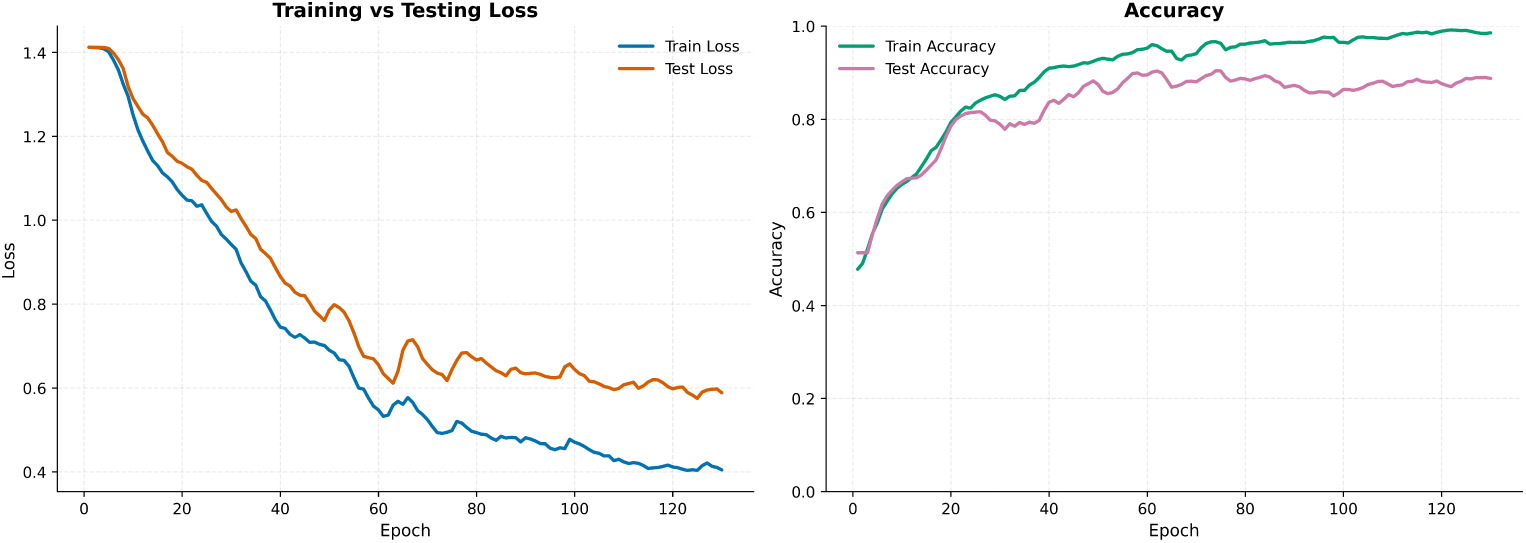
Performance of Triad-LMF on the BRCA 2 Dataset.

## 5 Discussion

In cancer research, multiomics data integration plays a crucial role in revealing the complexity of diseases and improving diagnostic Accuracy. Our proposed Triad-LMF model features a hierarchical fusion architecture that integrates progressively from local modality interactions to global information consolidation through the Two-Feature LMF module and Three-way LMF modules. This effectively captures the complex interactions among multiomics data, demonstrating promising application potential.

However, although the Triad-LMF model performs well overall, it was found that tri-modal fusion may introduce noise in specific tasks, leading to performance that is inferior to single-modal or dual-modal combinations, this may occur when modal redundancy occurs, or some modalities may carry more noise than signal. Including them can dilute the predictive power of strong modalities. This indicates that balancing information integration and noise control during the fusion process is a key challenge. In future work, we plan to investigate more adaptive fusion mechanisms that dynamically optimize modality selection and fusion weights[44], or Redundancy-adaptive multimodal learning[45] to mitigate the impact of redundant information, thereby enhancing model robustness and Accuracy. Additionally, given the complexity and diversity of cancer research, we intend to expand the model to incorporate more types of omics data. Considering the efficiency and generalizability of Triad-LMF, we believe that through these improvements and extensions, the model will play a more critical role in pancancer subtype classification, biomarker discovery, and precision medicine research, providing a more powerful tool to advance multiomics-driven cancer studies and accelerate the individualization and precision of cancer diagnosis and treatment. Moreover, in the absence of considering label noises, which some subtypes might be mislabeled [46], future work could focus on developing robust methods against label noises.

## 6 Conclusion

This study proposes a deep learning framework called Triad-LMF based on low-rank multimodal fusion, aiming to address the challenges of integrating multiomics data for cancer subtype classification. Through extensive experiments on BRCA, STAD, and BRCA 2 datasets, we demonstrate the significant advantages of Triad-LMF in classification performance, robustness, and generalization ability. Experimental results show that Triad-LMF achieved an F1 score of 0.8613 on the BRCA dataset, 0.8330 on the STAD dataset, and an Accuracy of 0.9237 on the BRCA 2 dataset, surpassing traditional machine learning baselines and advanced multiomics fusion methods such as MOGONET. UMAP visualization validated the critical role of Global Triadic Fusion in enhancing the latent space’s ability to distinguish subtypes, manifested by more compact subtype clusters and clearer boundaries between clusters. Ablation studies further confirmed the necessity of both Local Pairwise Fusion and Global Triadic Fusion; removing either component led to performance degradation, highlighting the synergistic effect of the hierarchical fusion architecture. SHAP-based feature importance analysis

These findings provide important evidence for identifying subtype-specific biomarkers and developing targeted therapeutic strategies.

## Ethics approval and consent to participate

Not applicable

## Consent for publication

Not applicable

## Availability of data and material

The multiomics datasets from The Cancer Genome Atlas (TCGA) are publicly available. The original contributions presented are available upon further inquiry and can be directed to the corresponding authors.

## Competing interests

The authors declare that the research was conducted in the absence of any commercial or financial relationships that could be construed as a potential conflict of interest.

## Funding

This work was supported by the National Natural Science Foundation of China under Grants (62276034, 62306052), Group Building Scientific Innovation Project for Universities in Chongqing (CXQT21021), Joint Training Base Construction Project for Graduate Students in Chongqing (JDLHPYJD2021016), and Science and Technology Research Program of Chongqing Municipal Education Commission (KJQN202100712)

## Author Contributions

XC and LZ conceptualized the project, XT and XC wrote and edited the original manuscript. XT and XC performed coding. XT conducted experiments. RT visualized the results. MJ and QC curated the data. DY and LZ edited the manuscript, LZ supervised the study. All authors read and approved the final manuscript.

## Acknowledgments

The authors would also like to thank Mr. Xiongtao Xiao for valuable discussion and thoughtful comments on the manuscript and language revision.

## Notes

### Competing Interest Statement

The authors have declared no competing interest.

### Summary of Updates

Changed to revised version, with extra analysis and correct writings

